# Emergent *Salmonella enterica* serovar Infantis forms a monophyletic lineage shaped by geographic structuring

**DOI:** 10.1101/2025.08.25.672257

**Authors:** Alejandro Piña-Iturbe, Daniel Tichy-Navarro, Josefina Miranda-Riveros, María José Navarrete, Andrea I. Moreno-Switt

**Affiliations:** Escuela de Medicina Veterinaria; Facultad de Agronomía y Sistemas Naturales, Facultad de Ciencias Biológicas y Facultad de Medicina; Pontificia Universidad Católica de Chile; Santiago; Chile

## Abstract

Multidrug-resistant *Salmonella* Infantis carrying pESI-like megaplasmids have disseminated worldwide representing a serious threat to public health. Previous studies have investigated its population structure and temporal dynamics above the continental level. However, their conclusions were constrained by limited datasets and sampling biases. To address these issues, we analyzed all publicly available *Salmonella* Infantis genomes to characterize its global population structure and phylogeographic dispersal. We selected a non-redundant dataset of 14,012 genomes representing the temporal, geographic, isolation source, and genomic diversity of *Salmonella* Infantis from 78 countries across five continents, collected between 1910 to 2024. Phylogenomic analyses showed that emergent megaplasmid-positive *Salmonella* Infantis forms a monophyletic lineage with significant geographic structuring. The megaplasmid-positive lineage was inferred to be originated in West Asia around 1990, followed by multiple introductions into Europe and a single transmission to South America which resulted in the dissemination of this pathogen to Northern America, and from there to the rest of the continent. Multiple recent transmission events of the American lineage to all continents were observed, driving the dispersal of the *bla*_CTX-M-65_ gene encoding extended-spectrum β-lactamases. Moreover, genomic evidence also suggests that the emergence of ESBL-producing strains in parts of Asia and Africa may be associated to poultry trading from the Americas. Our findings underscore the urgent need for integrating global human, animal, and environmental surveillance data with population genomic analyses to contain the threats posed by ESBL-producing *Salmonella* Infantis.

## INTRODUCTION

Non-typhoidal *Salmonella* (NTS) is one of the major causes of foodborne illness worldwide, producing millions of cases and thousands of deaths each year (1,2). NTS infection usually presents as a self-limited gastroenteritis, however, children under 5 years, immunocompromised people and the elderly, are at a higher risk of developing a severe invasive form of the infection (3). Moreover, the acquisition and dissemination of antimicrobial resistance (AMR) in NTS, especially against fluoroquinolones and third-generation cephalosporins (3GC), made this pathogen a global threat of high priority in the World Health Organization Bacterial Priority Pathogens List (4).

While NTS comprises more than 2,600 serovars, most cases of human infection are produced for a few highly prevalent serovars such as *Salmonella* Enteritidis and *Salmonella* Typhimurium (5,6). Nonetheless, multidrug-resistant *Salmonella enterica* serovar Infantis has emerged and disseminated globally in the last two decades, even displacing the historically most prevalent serovars in some geographic areas (7–11). Emergent *Salmonella* Infantis has been associated mainly with poultry and poultry-derived products (12). However, recent studies show that it is also disseminated in other animal and environmental niches including wild birds and river/irrigation waters (11,13,14). The success of emergent *Salmonella* Infantis is linked to the acquisition of a ≈300 kbp conjugative pESI megaplasmid and its variants (pESI-like megaplasmids), which encode several factors favoring the bacterium capacity of biofilm production, adhesion, invasion, and resistance to multiple antibiotic classes, including aminoglycosides (e.g., gentamycin), folate pathway inhibitors (e.g., trimethoprim-sulfamethoxazole; SXT) and 3GC (e.g., cefotaxime) with an extended-spectrum β-lactamase (ESBL) phenotype (11,15–17). In addition, emergent *Salmonella* Infantis also carries chromosomal mutations in the quinolone resistance-determining region (QRDR) of the *gyrA* gene, which result in intermediate resistance to fluoroquinolones and, in combination with plasmid-borne quinolone-resistance genes (e.g., *qnrB19*) produces a resistant phenotype (11). Resistance to SXT, 3GC and fluoroquinolones greatly limits the first line treatment options for severe infections caused by emergent *Salmonella* Infantis which has become a serious global concern (3,12).

Despite the importance of emergent *Salmonella* Infantis for human health and the spread of antibiotic resistance genes, there is still a limited understanding of its global population structure, geographic distribution of linages, and dissemination dynamics. Most studies addressing these topics are limited to the country or regional scale and base their conclusions on a few hundred isolates. Nevertheless, the current evidence shows that different subpopulations of emergent *Salmonella* Infantis may be associated with certain megaplasmid variants (8,17), ESBL-gene variants (9,18–20), *gyrA* mutations (21), and geographic locations (21,22).

The most comprehensive study to date, recently published in 2024, analyzed 5,284 *Salmonella* Infantis isolates obtained from 1989 to 2019, representing 74 countries (22). Bayesian clustering of 1,288 genomes representative of the total dataset identified three main clusters (A, B, and C). Emergent *Salmonella* Infantis clustered within clusters B (Global cluster) and C (European cluster). While cluster B included both pESI-positive and -negative isolates, cluster C was composed of pESI-positive isolates only. A core-SNP phylogenomic tree showed cluster B as paraphyletic and holds a pESI-negative clade diverging from the pESI-positive isolates. Although the authors did not estimate the time for megaplasmid acquisition by emergent *Salmonella* Infantis, their data suggest it was before 1987, the estimated time of the most recent common ancestor for cluster C (22).

Only one other study, also published in 2024, examined the structure of *Salmonella* Infantis at a global level, including only 415 isolates obtained between 1985 and 2019 from 15 countries, where 207 came from England and Wales (21). In this work, the core-SNP phylogeny placed all pESI-positive isolates as a paraphyletic group comprised of three Bayesian hierarchical clusters (clusters 1, 3 and 4) where pESI-negative isolates diverged from the pESI-positive ones. Conversely, a time-measured maximum clade credibility tree showed the pESI-positive isolates diverging from the negative ones around 1987 and suggested a possible origin of emergent *Salmonella* Infantis in Europe (21). This study showed a geographic clustering of the pESI-positive isolates originating from American (USA, Ecuador, Perú) and several European countries in clusters 1 and 3 which correlated with the presence of the *gyrA* D87Y and S83Y mutations, respectively (21). The authors also observed HC20 clusters (isolates linked by 20 or less cgMLST allele differences) comprised exclusively by pESI-positive strains (clusters HC20_343, 1369, 1272, and 7828) but no correlation or association with their geographic origin was explored.

To date, there are no studies that provide a global analysis of the geographic dispersal of emergent *Salmonella* Infantis. Therefore, a comprehensive study of megaplasmid-positive *Salmonella* Infantis phylogeography may contribute to a clearer understanding of its origin and the factors involved in its successful dissemination across the globe. In this study, we analyzed 26,981 *Salmonella* Infantis genomes isolated from 1985 to 2024 in 78 countries, representing 5 continents, unveiling that *Salmonella* Infantis carrying pESI-like megaplasmids form a monophyletic group comprised by geographically associated subpopulations. Importantly, Bayesian inference of the phylogeographic dispersal of emergent *Salmonella* Infantis indicated its origin on Western Asia and revealed the global dissemination of an ESBL-encoding lineage from the Americas.

## METHODS

### Genomes dataset

On July 12^th^, 2024, the Enterobase (23) database was queried for all *Salmonella* Infantis genomes according to serovar prediction by SISTR1. The 26,981 genomes were filtered by quality (N50 ≥ 100,000; coverage ≥ 30X), ambiguous serovar prediction (more than one SISTR1 serovar), and to eliminate redundant genomes based on geographic, temporal, niche, and genomic information; i.e., genomes linked by ≤5 cgMLST allele differences (see Population structure methods below) obtained from the same country, year, and isolation source were excluded and one representative genome was selected. This selection process resulted in 14,012 genomes whose assemblies were downloaded to the local storage.

### Identification of antibiotic-resistance genes/mutations and presence of pESI-like megaplasmids

AMR genes/mutations were identified on the downloaded assemblies using AMRFinderPlus v.3.12.8 with its database version 2024-07-22.1 (24). The pESI-like megaplasmids were identified using ABRicate v1.0.1 (https://github.com/tseemann/abricate) with default parameters, and a custom database comprising the 315 CDS from megaplasmid pN55391 (GenBank accession NZ_CP016411.1) (8), as previously described (11). The number of pN55391 genes in each of the 14,012 genomes was recorded together with the presence of complete megaplasmid backbone regions. To account for megaplasmid plasticity and potential missing genes, each one of the eight backbone regions (25) was considered complete if: all two genes were present for the toxin/antitoxin systems *pemK/I*, *vapB/C* and *ccdA/B*; there was at most one absent gene for the K88-like and Ipf fimbria gene clusters, the yersiniabactin synthesis cluster, the mercury resistance cluster (4 to 11 total genes in these regions); and, there were at most 5 out of 50 missing genes (10%) for the conjugative transfer region. As determining pESI-like megaplasmids presence based solely on presence of the backbone regions excluded genomes with high number of megaplasmid genes (**Fig. S1**), we decide to include the gene-number variable for megaplasmid presence assessment. Therefore, a genome was considered positive for a pESI-like megaplasmid if it harbored 7 out of the 8 backbone regions and/or 220 megaplasmid genes. A detailed description of the 220-gene cut-off is provided in the **Supplemental Material** and **Figs. S2, S3**.

### Population structure and phylogenomic analyses

The genome dataset was analyzed in Enterobase to determine the 7-gene MLST (based on genes *aroC*, *dnaN*, *hemD*, *hisD*, *purE*, *sucA* and *thrA*), cgMLST (cgMLST v2 based on the *Salmonella* 3,002 allele scheme; https://pubmlst.org/bigsdb?db=pubmlst_salmonella_seqdef&page=schemeInfo&scheme_id=4), hierarchical clustering of cgMLST profiles (HierCC v1), and minimum spanning trees (MSTs; MSTree v2).

For further analyses, the hierarchical cluster grouping all the pESI-positive genomes was selected (HC200_36; 13,833) and filtered to exclude genomes without information regarding collection year or country of isolation and to retain most of the temporal, geographic, genomic diversity and megaplasmid presence information. Therefore, one representative genome was selected for those isolates grouped in the same HC50 cluster (linked by ≤50 cgMLST allele differences), obtained the same year from the same country and with the same megaplasmid presence/absence status. The selected 821 genomes were used to construct a genome-wide multiple sequence alignment with Snippy v4.6.0 https://github.com/tseemann/snippy) using the closed chromosome of *Salmonella* Infantis strain N55391 (Enterobase assembly barcode SAL_XA0374AA_AS) as the reference.

The genome-wide alignment was uploaded to the Chilean National Laboratory for High Performance Computing (NLHPC) cluster where recombination events were detected with Gubbins v3.3 (26). Five iterations of the analysis were carried out (default setting) in which the first tree was built with IQ-Tree Fast (27) using the Jukes-Cantor (JC) nucleotide substitution model. Then, the trees in the remaining iterations were built with IQ-Tree Fast with the best-fitting substitution model as selected by ModelFinder (28) and IQ-Tree within Gubbins. The final selected model was the General Time Reversible model with a discrete Gamma model with four rate categories, empirically determined base frequencies and ascertainment bias correction (GTR+G4+F+ASC). Then, the output file containing the core-SNP alignment and masked recombination regions serve as the input for SNP-sites v2.5.2 (29) to obtain a recombination-free core-SNP alignment.

A recombination-free maximum likelihood core-SNP phylogenomic tree was built with IQ-Tree MPI multicore v2.3.2 in the NLHPC using the GTR+G4+F+ASC model (27). Tree topology support was estimated from 1,000 ultrafast bootstraps [UFBoot; (30)]. The phylogenomic tree was uploaded to iTOL v7 (https://itol.embl.de/) and rooted according to the temporal root (best-fitting root) estimated with TempEst v1.5.3 (31) using the collection year data and optimizing for the correlation coefficient.

### Temporal analysis

Genomes from clade IV of the phylogenomic tree were selected (441 genomes, including the N55391 reference) and a recombination-free core-SNP multiple sequence alignment was built as described above. A maximum likelihood phylogenomic tree was built with IQ-Tree MPI (GTR+G4+F+ASC) and then examined with TempEst, estimating the best fitting root maximizing the correlation coefficient. The analysis showed a linear trend (r=0.5415) with notable scatter, indicating a temporal signal and that a relaxed molecular clock would be appropriate for this data. After examination of the root-to-tip and residuals plots, seven genomes were identified as outliers and were excluded from further analyses (**Figs. S4, S5**).

A new recombination-free core-SNP multiple sequence alignment of the megaplasmid-positive clade was built (307 taxa; 7,903 SNPs). This alignment was the input for construction of the time-measured phylogeny using BEAUti v1.10.4 and BEAST v.1.10.4 (32). The nucleotide substitution model selected with ModelFinder/IQ-Tree in Gubbins (GTR+G4+F) was also applied here. As no constant sites are present in a SNP alignment, ascertainment bias correction is needed to avoid artifacts on temporal calibration of the trees (33). Ascertainment bias correction was made explicit in the BEAST XML file by manually specifying the number of constant nucleotide bases (33) (A: 1123516, C: 1229301, G: 1237516, T: 1128879; see the **Supplemental Material** for the calculation details). The uncorrelated lognormal clock (UCLN) was selected over the strict clock since a preliminary test (30M states, 5M burn-in) showed that the distribution of the rate variation coefficients of the UCLN clock excluded the zero value, providing evidence against a constant rate across tree branches (strict clock). To assess the best population model, the selected nucleotide and clock models were combined with the Exponential Growth, Logistic, or Bayesian Skyline (7 groups, skyline model: piecewise-linear) models. The clock rate (ucld.mean) and initial population size (pop.Size) parameters were sampled from lognormal distributions with the following parameters: ucld.mean – initial value: 1.0, mean in real space: 1.9E-7 (34), standard deviation in real space: 2.0; pop.Size – initial value: 1.0, μ=10, σ=2.0 (21). For the Logistic model, the logistic.t50 parameter was sampled from a uniform distribution (lower value: 0.0, upper value: 100.0). For the Skyline model, the pop.Size prior distribution was kept as default. To improve mixing, the weight for the clock operators (ucld.mean, ucld.stdev) were set from 3.0 to 5.0. The MCMC chain was set to 200M, sampled every 5000 states. The Bayes factors (log_10_) for each model combination were calculated from the log marginal likelihoods estimated using the path sampling/stepping-stone sampling in three independent MCMC simulations (chain length 2M with 100 path steps). All simulations were run at the NLHPC. The Bayesian Skyline model was finally selected over the Exponential Growth and Logistic models.

Convergence of the MCMC chains and effective sample size (ESS) of the different parameters was assessed with Tracer v1.7.2 (35). To reach an ESS higher than 200, the independent simulations and trees were combined with LogCombiner v1.10.4 (https://beast.community/logcombiner). Finally, a time-measured maximum clade credibility (MCC) tree was generated with TreeAnnotator v1.10.4 (https://beast.community/treeannotator).

### Phylogeographic analysis

To reconstruct the geographic transmission events, the country of isolation for each genome in the megaplasmid-positive clade was classified according to the United Nations standard M49 and grouped by subregion (36). The subregion information for each genome was included in BEAUti as a new partition and the strict clock was selected for it. The ancestral states were reconstructed for the subregion partition. The best discrete substitution model was selected between the Symmetric and Asymmetric models (with Bayesian Stochastic Search Variable Selection, BSSVS) according to the log_10_ Bayes factor, as described in the previous subsection. The Asymmetric model was selected, and the inferred phylogeographic dispersal of emergent *Salmonella* Infantis was visualized in the MCC tree with iTOL, and in the world map using spread.gl (37). The geographic centroids for each subregion were used to map their location, and only significant transmissions are depicted, based on log_10_(Bayes factor) > 3, as estimated through BSSVS.

### Statistical analysis

Person’s χ^2^test was used to assess whether HC20 cluster and geographic (continent) variables were independent. The chisq.test function was used in RStudio v2023.06.1 (R v4.4.3) to assess a contingency table containing the number of genomes present in clusters HC20_343, HC20_7828, HC20_7264, and HC20_7377 per continent (Americas and Europe). As the contingency table contained several zeroes or <5 values, a Monte-Carlo simulation with 100,000 replicates was specified. The residuals were plotted and values ≥2 or ≤-2 were considered over and underrepresented, respectively. GraphPad Prism v10.5.0 was used to run a Mann-Whitney test to compare the megaplasmid-gene distributions from genomes within and outside the pESI+ subclade. Fitch-parsimony score permutation test was used to test phylogeographic structuring within the pESI+ subclade. The parsimony score was calculated using a custom Biopython script and was compared with the resulting distribution of scores resulting from 100,000 random tip–trait assignments. For all statistical analyses the α value was set to 0.05.

## RESULTS

To study the global population of *Salmonella* Infantis, 14,012 genomes representing the geographic, niche, temporal, and genomic diversity of this pathogen were analyzed (**Table S1, S2**). While the geographic component of the dataset is biased towards the countries performing most of the *Salmonella* sequencing globally (United States: 8,595; United Kingdom: 1,846; France: 446; Canada; 203), the dataset represents 78 countries from 5 continents, and 14 countries are represented by more than 100 genomes (**Fig. 1A**). The genomes came from isolates obtained from animal, environmental, food and human niches, mostly from poultry (6,131) and clinical human origin (3,138) (**Fig. 1B**). The dataset contains genomes from isolates obtained from 1910 to 2024. However, the majority (12,940/12,948) were isolated after 1992, with numbers increasing exponentially until 2019. The oldest emergent *Salmonella* Infantis genomes (positive for pESI-like megaplasmid) derived from human-infection isolates obtained in years 1999 (4 genomes) and 2000 (18 genomes) in Japan (38) (**Fig. 1C**). We used this dataset to explore the global population structure and dynamics of emergent *Salmonella* Infantis.

**Figure 1.**
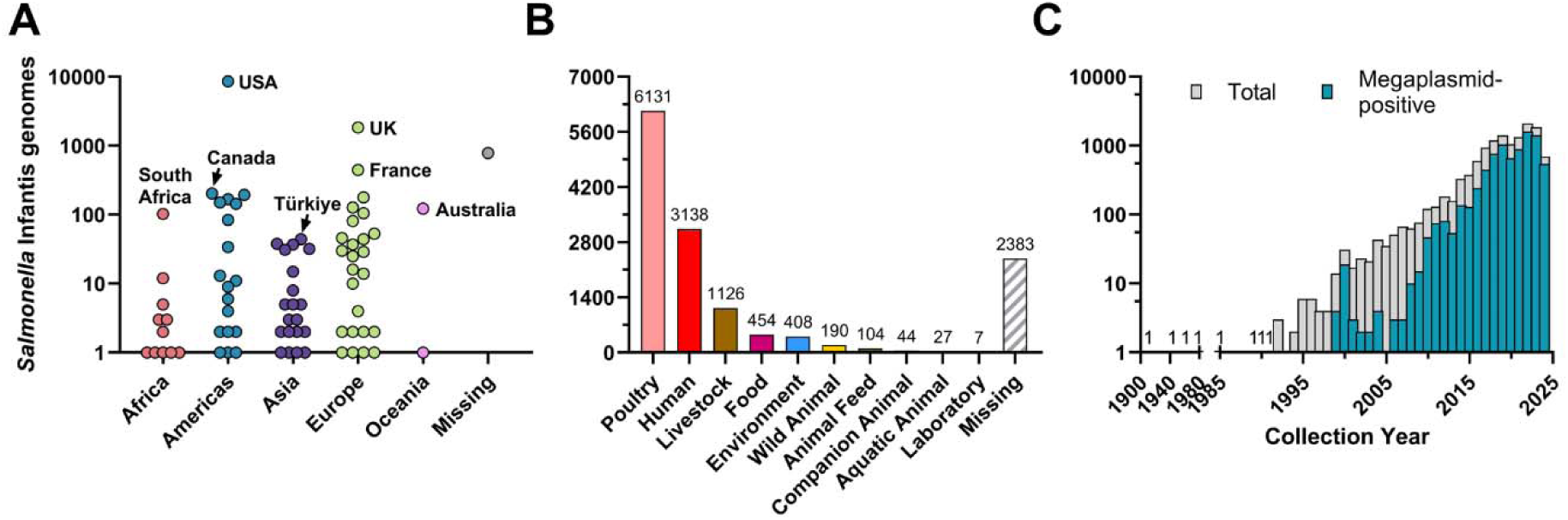
Geographic location, source niche, and collection year of the global *Salmonella* Infantis genome dataset. Number of *Salmonella* Infantis genomes **A)** per country of isolation and colored by United Nations’ geographic region (continent); **B)** per isolation source according to the Enterobase classification; and per year of isolation. In A, the countries contributing with most of the dataset genomes are indicated. In C, the number 1 indicates the presence of 1 genome, and the genomes containing pESI-like megaplasmids are colored in cyan-blue.

### Emergent *Salmonella* Infantis belongs to predominant HC20 clusters showing geographical association

Hierarchical clustering of genomes linked by ≤20 cgMLST alleles (HC20 clusters) resulted in 1,727 clusters. However, 52.5% (7,358/14,012) of the genomes were grouped within clusters HC20_343 and HC20_775 (**Fig. 2A, D**). Importantly, we observed an uneven geographic distribution of clusters in which almost all genomes from a particular HC20 cluster came from the same continent, such as HC20_220954 from Africa (100%; 10/10), HC20_343 from the Americas (97.6%; 6,113/6,263), HC20_136276 from Asia (87.1%; 27/31), HC20_7828 from Europe (97.9%; 285/291), and HC20_45736 from Oceania (96.5%; 55/57), among others (**Fig. 2B, 2E; Table S3**). When the presence of pESI-like megaplasmids was assessed, the minimum spanning tree showed two clearly delimited groups of positive and negative genomes (**Fig. 2C**). Noteworthy, these pESI-like-positive isolates were almost exclusively grouped in specific HC20 clusters, being HC20_343, HC20_7828, HC20_7264, and HC20_7377 the most representative (**Fig. 2F**). Interestingly, while clusters HC20_7828, HC20_7264, and HC20_7377 were predominant in European countries, the cluster HC20_343 had a broader distribution —being present in all continents— and comprising almost all isolates from the Americas, and the majority from Eastern Asia, Southeastern Asia, Australia and New Zealand, and Southern Africa (**Fig. 3; see also Fig. 2E**). Statistical testing for independence between the HC20 predominant clusters and the Americas and Europe continents indicated a statistically significant (Person’s χ^2^; p=0.00001) association with cluster HC20_343 being overrepresented in the Americas, and clusters HC20_7828, HC20_7264, and HC20_7377 being overrepresented in Europe (**Supplemental Material**). Together, our analysis reveals the association of pESI-like megaplasmids with particular *Salmonella* Infantis HC20 clusters and of these clusters with geographic subregions.

**Figure 2.**
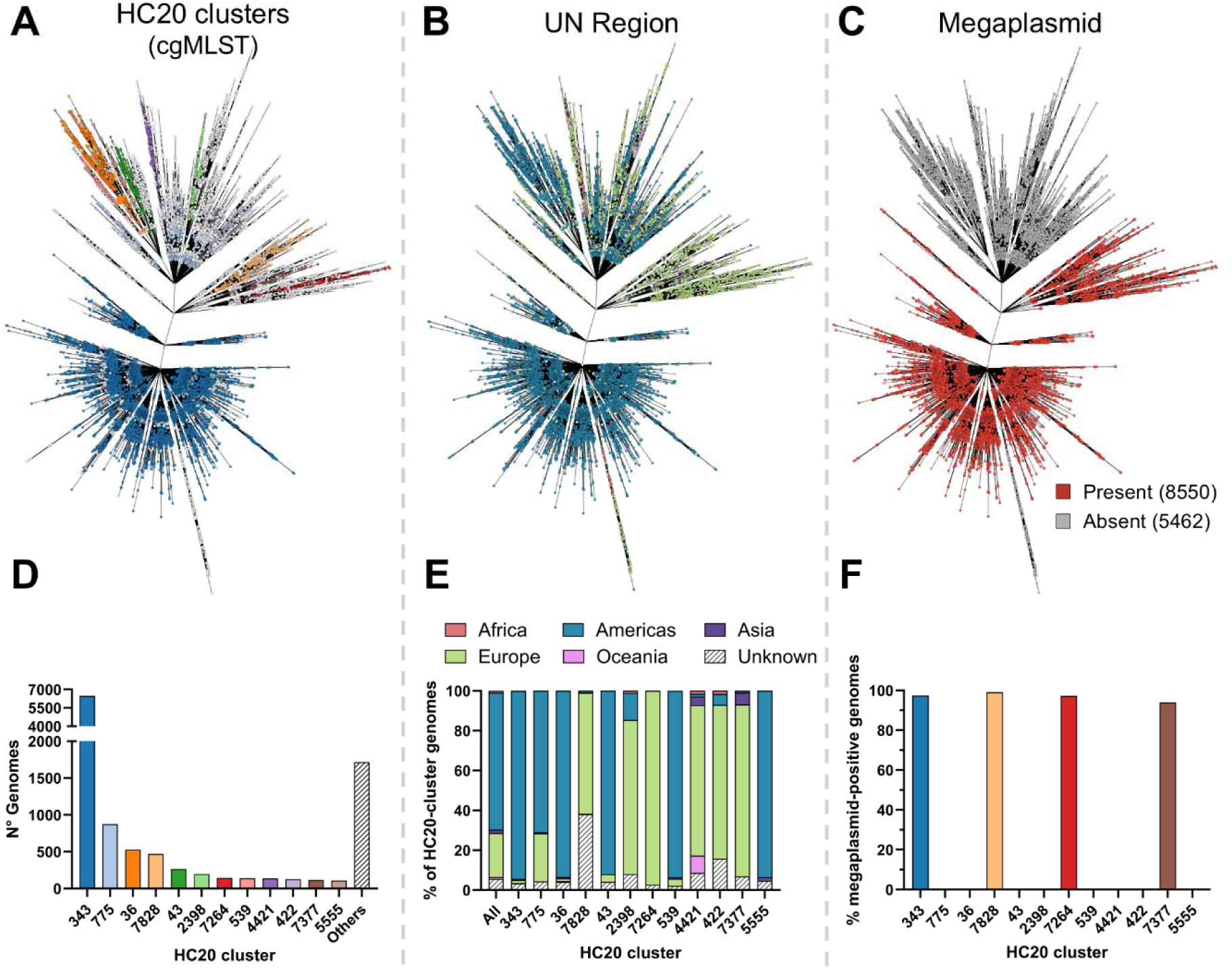
Global population structure of *Salmonella* Infantis. Minimum spanning trees (MSTs) depicting the population structure of 14,012 genomes of *Salmonella* Infantis from 78 countries. Nodes are colored according to **A)** HC20 cluster, **B)** United Nations’ continental region, and **C)** megaplasmid presence. The branch lengths are in logarithmic scale to improve visualization. Below the MSTs, the bar charts show the *Salmonella* Infantis genomes grouped by HC20 cluster and **C)** showing their absolute frequency, **E)** relative frequency across the UN continental regions, and **F)** relative frequency of megaplasmid-positive genomes. Only the HC20 clusters with 100 or more genomes are shown.

**Figure 3.**
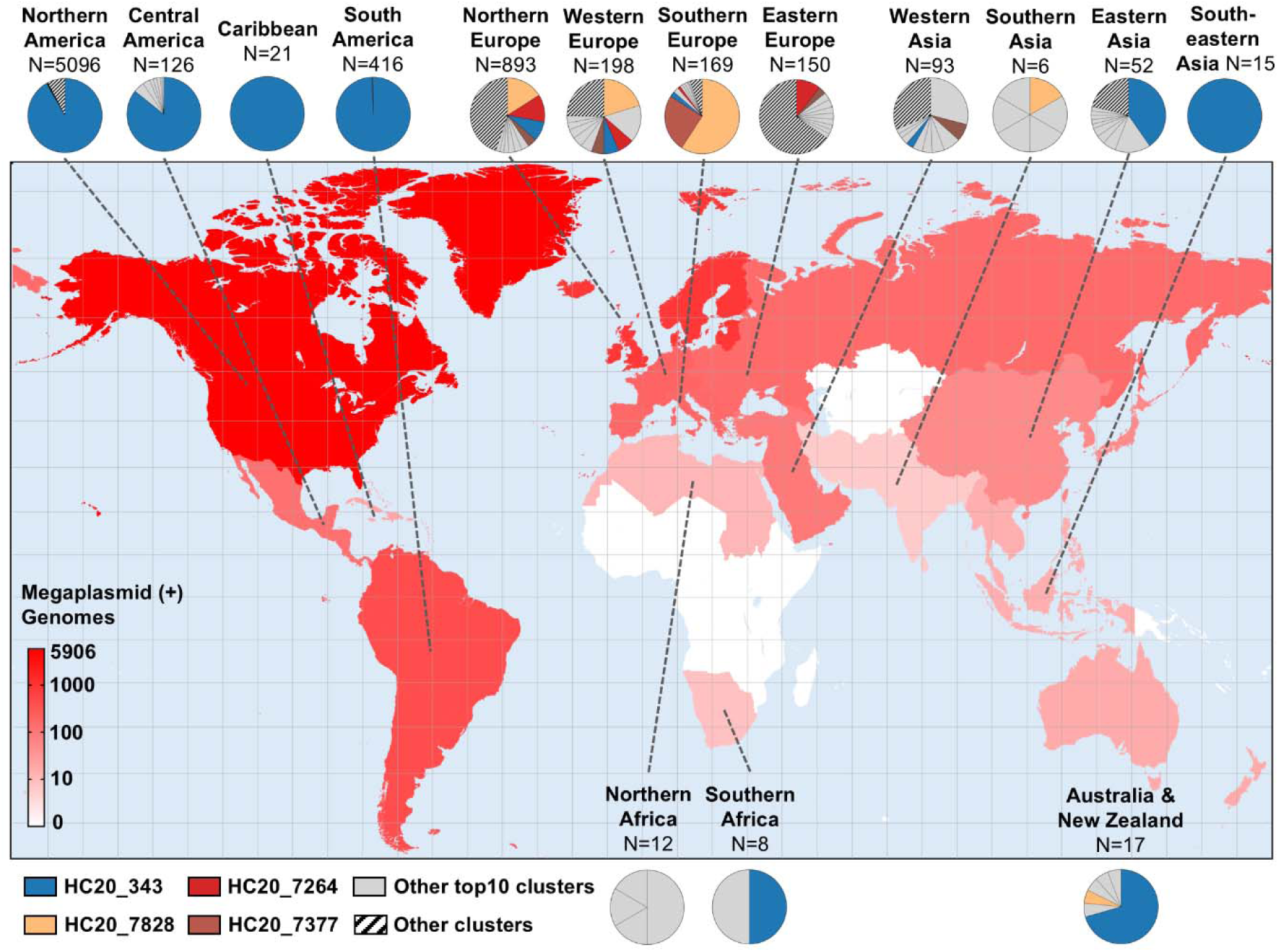
Geographic distribution of emergent *Salmonella* Infantis HC20 clusters. A Gall stereographic projection of the world subdivided in the 22 United Nations’ subregions and colored to show the number of megaplasmid-positive *Salmonella* Infantis genomes per subregion according to the color scale in the lower-left corner (total: 8,083 genomes with geographic metadata). Antarctica was excluded from the map to improve visualization. The pie charts show the relative distribution of the top 10 most frequent megaplasmid-positive HC20 clusters per geographic subregion, but only the most frequent clusters at the world level are highlighted (HC20_343, HC20_7828, HC20_7264 and HC20_7377) to show their global distribution. The remaining top10 HC20 clusters per subregion are colored in gray, and those clusters outside the top10 are colored in white with black diagonal lines. The Gall stereographic projection was made and colored in https://www.mapchart.net/.

### Megaplasmid-positive *Salmonella* Infantis form a monophyletic group with geographically associated lineages

To assess the phylogenetic relationships of megaplasmid-positive *Salmonella* Infantis, we selected representative genomes belonging to the lowest hierarchical cluster encompassing all megaplasmid-positive isolates. Cluster HC200_36 grouped all (8,549/8,550) megaplasmid-positive genomes except by one for which cgMLST profiles could not be determined (**Table S1**). This cluster also included 5,284 megaplasmid-negative genomes, thus encompassing 98.7% (13,833/14,012) from the global dataset. Eight hundred twenty-one genomes — representative of clusters linked by ≤50 allele differences— were selected, retaining the geographic, temporal and megaplasmid-presence information (See Methods). The representative subset spanned 70 countries on five continents, with the largest contributions being from the United Kingdom (n=149), United States (n=84) and Germany (n=49), covering sample collection years from 1978 to 2024 (median = 2017; **Supplemental Material**).

We generated a recombination-free core-SNP maximum likelihood phylogeny rooted on the best-fitting temporal root (**Fig. 4**). This tree resolved the 821 HC200_36 genomes into four fully supported clades (UFBoot = 100%): Clade I (n = 98), Clade II (n = 130), Clade III (n = 153) and Clade IV (n = 440). All 293 megaplasmid-positive genomes meeting our presence thresholds (≥7 backbone regions or ≥220 megaplasmid genes) were grouped in a single maximally supported subclade within Clade IV, showing the megaplasmid-positive lineage as a monophyletic group that we termed the pESI+ subclade. Of note, alongside these 293 genomes, another 20 genomes (6.4%) labeled as megaplasmid-negative clustered within this subclade. However, they contained between 124 to 219 pESI-like megaplasmid genes indicative of false-negative assignations caused by threshold application and/or potential backbone-gene loss events. On the other hand, the remaining 508 megaplasmid-negative genomes located outside the pESI+ subclade harbored a significantly lower number of pESI-like megaplasmid genes (0 – 110 genes, p<0.0001; **Fig. S6**), further supporting the observation that pESI-like megaplasmids are present only in the identified subclade. Importantly, we observed a clear clustering of isolates by geographic origin within the pESI+ subclade, particularly for American and European genomes. A Fitch-parsimony score permutation test (100,000 permutations) resulted in a parsimony score of 41 which was significantly lower than expected under random tip–trait assignments (p = 0.00001), indicating strong phylogeographic structuring.

**Figure 4.**
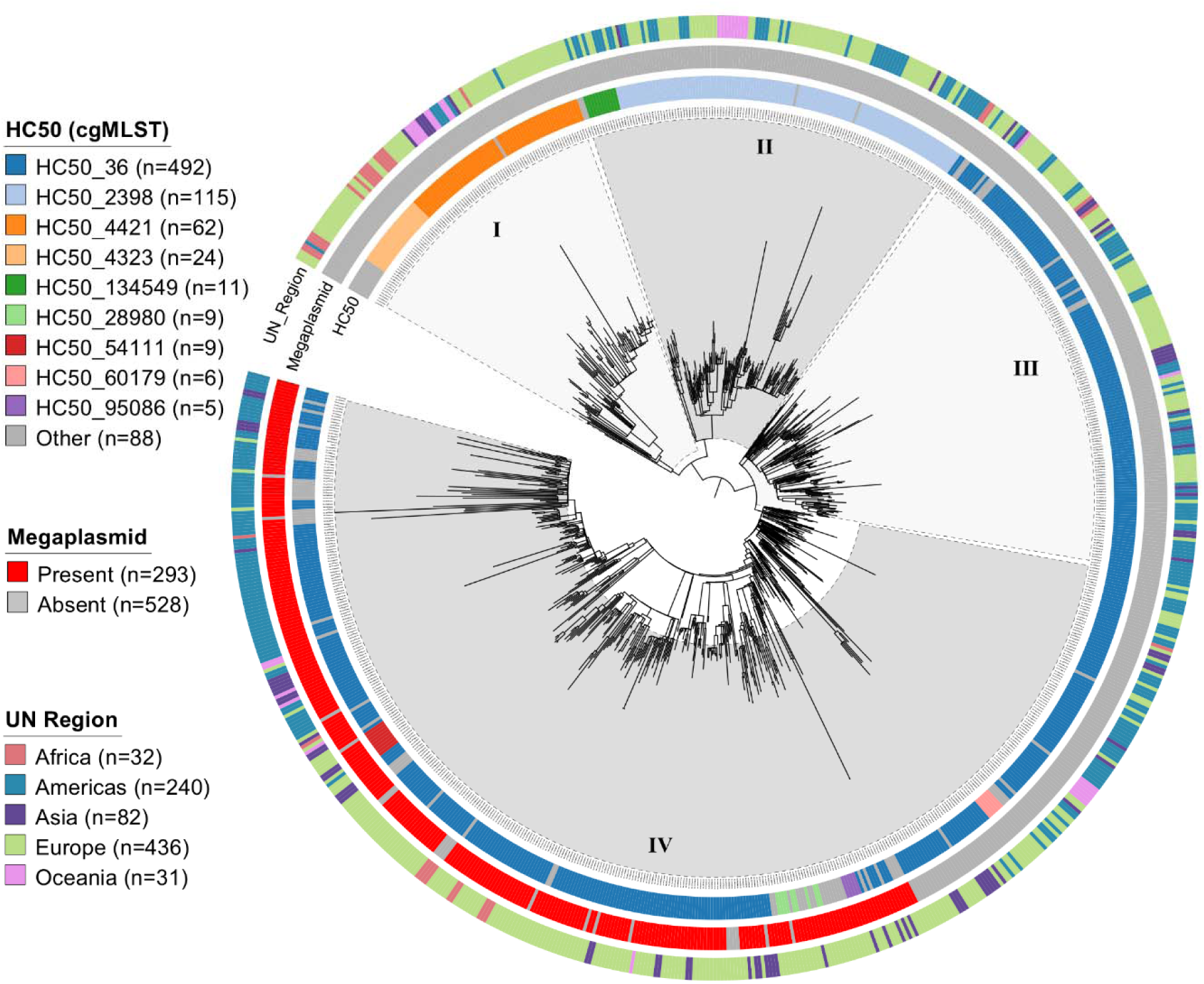
Phylogenomic tree of HC200_36 *Salmonella* Infantis. A maximum likelihood phylogenomic tree was built based on the recombination-free core-SNP alignment (17,718 sites) of 821 representative *Salmonella* Infantis genomes selected to retain most of the temporal, geographic, genomic diversity and megaplasmid-presence information within the HC200_36 cluster (see Methods for details). The phylogenomic tree was rooted according to the best-fitting temporal root estimated with TempEst. Major clades are indicated with roman numerals I-IV. The colored rings, from inside to outside, indicate the HC50 cluster, megaplasmid presence, and United Nations continental region for each genome.

Altogether, the phylogenomic and permutation-test analyses reveal the spread of a single emergent megaplasmid-positive *Salmonella* Infantis lineage, characterized by distinct geographic radiations.

### Dissemination of megaplasmid-positive *Salmonella* Infantis from the Americas is driving global spread of *bla*_CTX-M-65_-carrying strains

To assess the temporal and geographic transmission dynamics of megaplasmid-positive *Salmonella* Infantis we inferred a time-measured maximum clade credibility (MCC) tree (**Fig. 5**) and reconstructed the ancestral states for the geographic subregion variable (**Fig. 6**). According to these analyses, the most recent common ancestor of megaplasmid-positive *Salmonella* Infantis originated in Western Asia (subregion prob. 0.99) around 1990 (95% HPD: 1985.6-1993.6), followed by early transmission to Northern Europe around 1992 and expansion within this location (subregion prob. 0.94; node prob. 0.99). Most descendants from this clade harbor the S83Y mutation in the quinolone-resistance determining region (QRDR) of *gyrA*, suggesting that acquisition of this resistance mutation was an early event (**Fig 5**). The tree topology also indicates a well-supported (subregion prob. 0.90; node prob. 1.0) introduction to Eastern Europe around 1999, also succeeded by local expansion and further transmission to the remaining European subregions and Africa (**Figs. 5, 6A**). This clade is also characterized by the S83Y mutation in *gyrA*. After these events, a single introduction event was inferred from Western Asia to the Americas, specifically to South America around 2005 (95% HPD: 2002.5-2007.0), that lead to expansion of this lineage in the subregion and a rapid introduction to Northern America around 2009 (95% HPD: 2007.0-2010.5) and Central America around 2010 (95% HPD: 2007.9-2012.5) (**Figs. 5, 6A**). No other transmission events toward the Americas could be detected. From 2010 onwards, Europe and the Americas contributed with multiple transmission events of megaplasmid-positive *Salmonella* Infantis to the other continents in addition to local circulation and expansion of these lineages (**Fig. 6B**). Noteworthy, the phylogeographical dispersal of the American clade (subregion prob. 0.88; node prob. 1.0), characterized by the *gyrA* D87Y mutation and a high frequency of *bla*_CTX-M-65_-positive genomes, revealed transmission events to all other continents (except Antarctica) (**Fig. 5, 6**), evidencing the global dissemination of *Salmonella* Infantis harboring resistance determinants against quinolones and third-generation cephalosporines.

**Figure 5.**
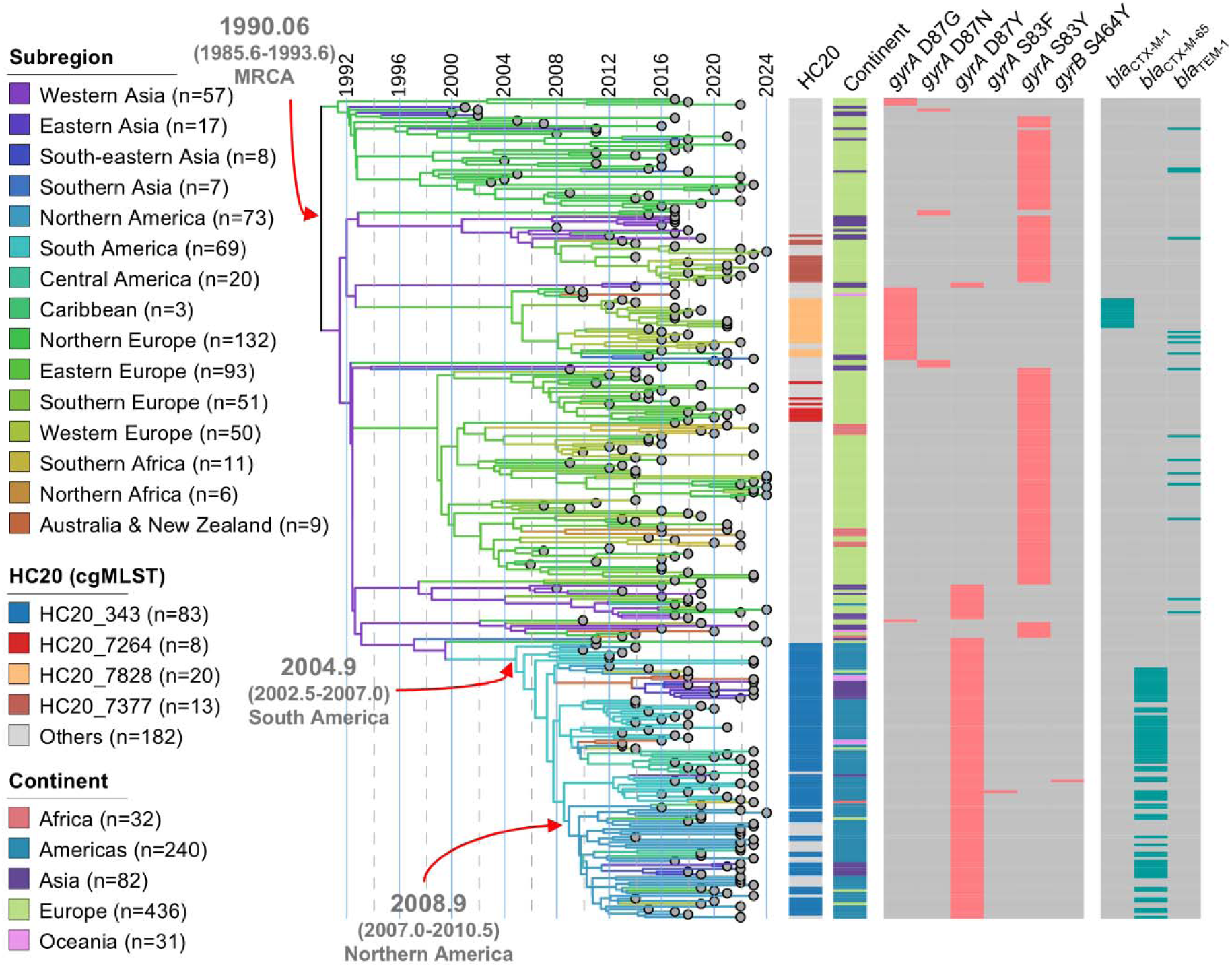
Time-measured phylogenomic tree of global emergent *Salmonella* Infantis. A time-calibrated maximum clade credibility tree inferred from 307 emergent *Salmonella* Infantis isolates (megaplasmid-positive clade in Fig. 4) representing 15 geographic subregions and 5 continents (Regions). Presence of quinolone resistance mutations in genes *gyrA* and *gyrB*, and β-lactam resistance genes are also indicated. The branches are colored according to reconstruction of ancestral states based on subregion. The median time to the most recent common ancestor (MRCA) for the emergent, South American, and North American clades are indicated together with their 95% high posterior density (HPD) values.

**Figure 6.**
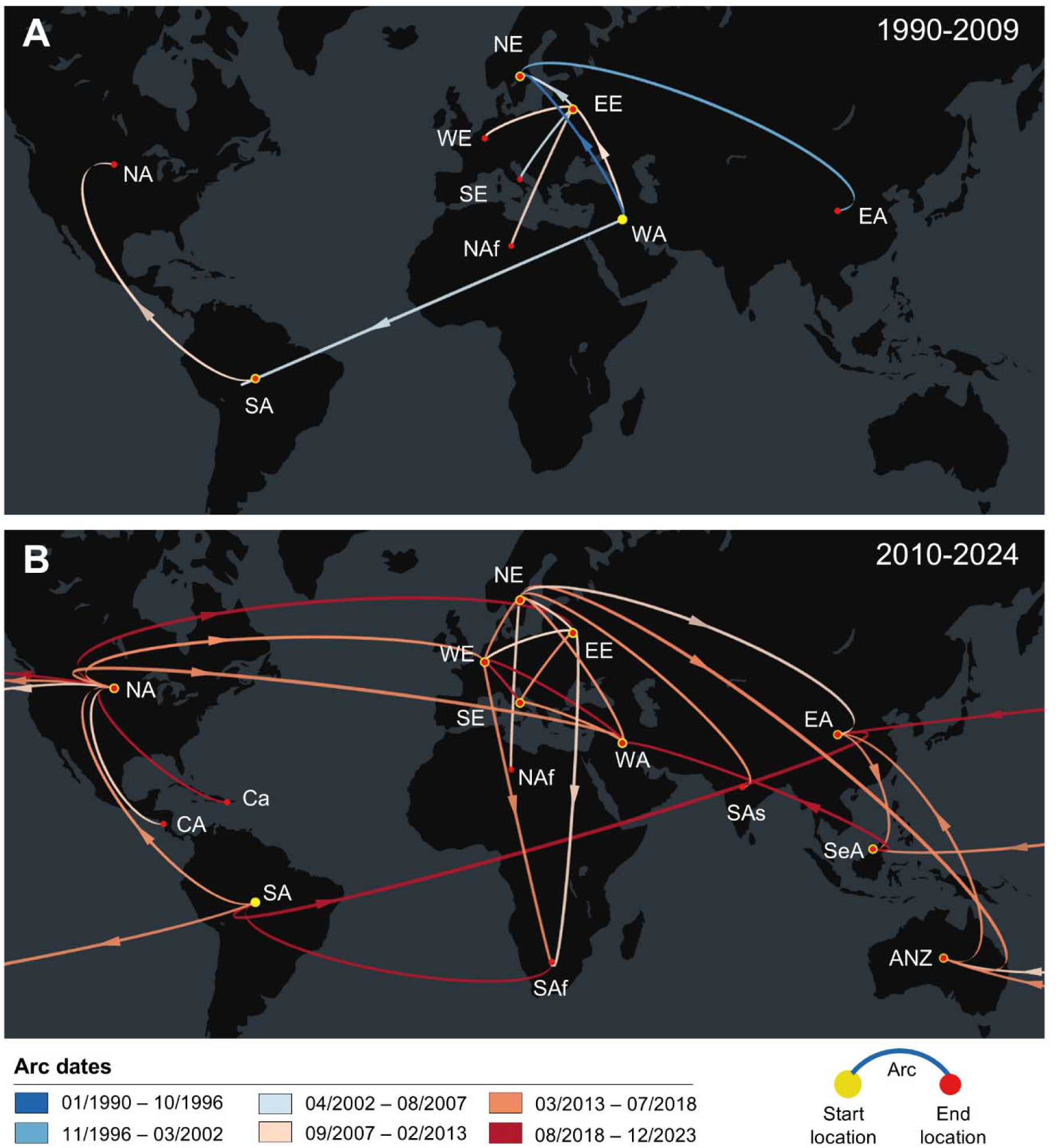
Phylogeographic reconstruction of the global dispersal of emergent *Salmonella* Infantis. Discrete spatiotemporal dispersal of emergent *Salmonella* Infantis from 1990 (emergence) to 2024, reconstructed using Bayesian inference. Panels show the dispersal before **(A)** and after **(B)** 2010. Arcs represent lineage dispersal between two geographic subregions, colored according to the inferred date (Arc dates). Points at both ends of the arcs indicate directionality: start point is yellow and the end point is red. Arrowheads were added to aid in visualizing directionality. Only significant transmissions are depicted, based on Log(Bayes factor) >3, as estimated through BSSVS. Abbreviations: NA: Northern America; CA: Central America; SA: South America; Ca: Caribbean; NE: Northern Europe; WE: Western Europe; SE: Southern Europe; EE: Eastern Europe; NAf: Northern Africa; SAf: Southern Africa; WA: Western Asia; SAs: Southern Asia; EA: Eastern Asia; SeA: South-eastern Asia; ANZ: Australia and New Zealand.

## DISCUSSION

*Salmonella* Infantis carrying pESI-like megaplasmids have emerged and disseminated worldwide in the last decades, representing a serious threat to public health, especially for people at higher risk, as this foodborne pathogen is associated with resistance against first-line antibiotics used to treat severe infections (12). A few previous studies have investigated the population structure and temporal dynamics of emergent *Salmonella* Infantis above the continental level (21,22). However, while providing valuable insights, the conclusions obtained and their generalization were limited by the size of the genome datasets analyzed, or by sampling biases that resulted in overrepresentation of genomes from particular sources or geographic locations. In addition, the approaches used in these works for bacterial clusters/lineage identification are dependent on the input alignment (39), resulting in cluster outputs that cannot be extrapolated outside the sample used. To address these issues, we decided to analyze all the *Salmonella* Infantis genomes available in Enterobase by July 12^th^ 2024, leveraging the hierarchical clustering of cgMLST profiles according to HierCC, which provide stable cluster identification and nomenclature (23,40), to assess the population structure and geographic associations among 14,012 genomes selected to retain most of the geographic, temporal, isolation source and genomic diversity information.

Our analyses revealed clear associations between *Salmonella* Infantis subpopulations, megaplasmid-presence, and geographic location. Out of >1,700 clusters linked by ≤20 cgMLST alleles, clusters HC20_343, HC20_7828, HC20_7264, and HC20_7377 were the most abundant among megaplasmid-positive *Salmonella* Infantis, grouping 82.1% (6,633/8,083) of the genomes. The HC20_7828, HC20_7264, and HC20_7377 clusters were associated with Europe and, in agreement with previous studies (9,21), the cluster HC20_7828 was found as the most abundant subpopulation within this continent, encompassing 20.1% (283/1,410) of the genomes. Overall, Europe displayed a high variety of megaplasmid-positive *Salmonella* Infantis, which was also reflected in the higher number of *gyrA* mutations identified, with D87G and S83Y being the most common, in agreement with previous reports (**Table S4**) (9). The observed predominance of cluster HC20_343 in the Americas aligns with previous studies describing the emergence and circulation of megaplasmid-positive *Salmonella* Infantis carrying specific features in different American countries, such as the ESBL-encoding *bla*_CTX-M-65_ gene and the *gyrA* D87Y mutation (11,19,41,42), which were carried by this cluster. We found the presence of other HC20 clusters in subregions from Africa, Asia and Oceania. However, despite the relatively low number of genomes available from these locations, the predominant American and European megaplasmid-positive clusters were also identified, underscoring the widespread distribution of these subpopulations across the globe.

Previous core-SNP analyses of *Salmonella* Infantis have produced phylogenomic trees with inconsistent topologies, either locating megaplasmid-negative genomes as diverging from the positive ones, or vice versa. Interpretation of these phylogenies was also limited by the reduced datasets used, sampling biases or the lack of methodological details such as how rooting was performed (17,21,22,43). We rooted our recombination-free core-SNP maximum likelihood (ML) tree using the best-fitted temporal root estimated with TempEst. This approximation resulted in a tree showing megaplasmid-positive *Salmonella* Infantis as diverging from megaplasmid-negative strains. This topology agrees with the published literature that reports the finding of megaplasmid-positive and/or MDR *Salmonella* Infantis in the last decade across multiple countries, indicating that the emergence of this pathogen is a relatively recent event (15,18,20,21,44–46). Importantly, the ML tree showed that the megaplasmid-positive *Salmonella* Infantis formed a monophyletic group. A previously published time-measured MCC tree inferred from 245 *Salmonella* Infantis genomes had a concordant topology (21). However, this tree mostly represented England and Wales, and few other countries from Europe and America. Conversely, the megaplasmid-positive clade our ML phylogeny (referred as the pESI+ clade in this work) encompassed 53 countries from 5 continents, therefore providing compelling evidence that the entire emerging lineage descends from a single common ancestor that acquired the megaplasmid, instead of multiple independent acquisitions of pESI-like megaplasmids by different *Salmonella* Infantis lineages.

Our phylogeographic reconstruction placed the MRCA of pESI+ *Salmonella* Infantis in Western Asia around 1990 (95% HPD 1985.6-1993.6) and identified a single introduction into the Americas through South America by 2005 (95% HPD 2002.5-2008.9), followed by rapid onward spread into North America around 2009 (95% HPD 2007.0-2010.5). This scenario is consistent with an early study of 160 genomes from the United States and Perú, which found a high probability (98.7%) that the latter was the origin of the United States isolates (19). Notably, despite differences in the number of genomes (160 vs. 307) and the geographic origins of isolates (2 vs. 53 countries) between the dataset analyzed in that earlier work and the present study, the tMRCA for the American isolates and its 95% HPD are almost coincident. We found that, for the Americas, Perú and the USA had the earliest megaplasmid-positive isolates, collected in 2009 and 2012 respectively (**Table S5**). Additionally, in our time-measured MCC tree, the Peruvian isolates were placed at the basal positions in the American clade alongside the earliest United States samples, further supporting Perú as an early hub of emergent *Salmonella* Infantis in the continent.

Carrying pESI-like megaplasmids confers significant advantages for the host bacteria, including increased biofilm production, tolerance to heavy metals and disinfectants, and resistance to multiple antibiotics, including third-generation cephalosporins in the isolates harboring *bla*_CTX-M_ alleles (15,18,19). These features have likely contributed to the extensive dissemination of emergent *Salmonella* Infantis in the global poultry industry, which is characterized by centralized sourcing of breeding stocks, and the dominance of a few global producers and exporters (25,47). As a consequence, poultry products, particularly broiler chicken and turkey, have been identified as the primary setting for megaplasmid-positive *Salmonella* Infantis amplification and the principal source of human infections (12,25). The analysis of the available isolation source metadata revealed two scenarios for the intercontinental spread of HC20_343 *Salmonella* Infantis isolates carrying the ESBL-encoding gene *bla*_CTX-M-65_ from the Americas. (**Fig. S7; Table S6**). When focusing on poultry and human sources, the HC20_343 isolates from Southern Africa (n=4), Western Asia (n=3) and Eastern Asia (n=10) were present in both niches, suggesting ongoing local circulation of this lineage in those poultry production systems. Accordingly, ESBL-producing *Salmonella* Infantis carrying the *bla*_CTX-M-65_ gene has been recently isolated from chicken farms and processing facilities in South Korea (48,49). We found that out of the 40 *Salmonella* Infantis available on Enterobase (Aug 25^th^, 2025), 33 were from the HC20_343 cluster and harbored *bla*_CTX-M-65_ (**Table S7**). Therefore, from the first scenario we hypothesize that the dispersal of the American HC20_343 *Salmonella* Infantis to Asia and Africa is mainly linked to the poultry industry. Conversely, in the second scenario, as observed in European subregions and Australia & New Zealand, poultry isolates were absent and most isolates with source data were from human clinical origin (90/96), followed by food (4/96) and environmental or animal origin (2/96). Although we do not have travel history information, a previous study has linked international travel to the Americas with HC20_343 *Salmonella* Infantis infections in Europe (9). Therefore, we hypothesize that human travel and/or contaminated broiler meat is playing the major role for the dispersal of *bla*_CTX-M-65_ to these locations, rather than local poultry reservoirs.

Our study has limitations that should be considered in interpreting the extent of our conclusions. First, public genome collections are geographically biased, and uneven source niche sampling can influence phylogeographic reconstructions. Therefore, we present the inferred origins and dispersal routes as hypotheses supported by our dataset rather than definitive events. In addition, to diminish the impact of these biases on our analysis, we selected representative genomes capturing geographic, temporal, source, and genomic diversity, filtering redundant genomes. While main genome contributors were the United States of America and the United Kingdom, 12 other countries across the five continents contributed with ≥100 genomes, making our study the most representative to date. Nevertheless, sequencing efforts in underrepresented regions are required to better depict the global population structure of emergent *Salmonella* Infantis and its worldwide dissemination. The high computational costs of temporal and phylogeographic analyses also posed an additional limitation. A greater number of character states (e.g., nucleotide at a given site or geographic location), longer MCMC chains to ensure sufficient effective sample sizes (>200), and the need to run independent replicate inferences to compare models and confirm convergence of posterior distributions all substantially increase the number of likelihood calculations —and, consequently, the overall runtime and required computing resources (32,50). As a consequence, performing phylogeographic analyses at the HC20 level or with country-level data was not possible. Finally, the limited available epidemiological metadata (e.g., missing isolation source, poultry/breeder-stock trading, and travel information) constrained our capacity to identify exact transmission pathways. Future work incorporating broader and balanced sampling for different sources, and integration of trading/epidemiological data will be needed to resolve remaining uncertainties.

### Concluding remarks

Current data links the infection with emergent *Salmonella* Infantis to higher hospitalization rates (6,19) and the presence of *bla*_CTX-M-65_ and the *gyrA* D87Y in HC20_343 isolates further limits treatment options for severe or invasive infections (3). Therefore, the predominance of HC20_343 *Salmonella* Infantis in the Americas and its dissemination to other continents underscores the urgent need for preventive measures and targeted interventions to control the spread of this pathogen. Moreover, multiple *Salmonella* serovars —such as Agona, Alachua, Muenchen, Senftenberg, and Schwarzengrund— have acquired *Salmonella* Infantis pESI-like megaplasmids, including those harboring the *bla*_CTX-M-65_ gene, revealing the introduction of critical resistance determinants into new bacterial hosts (25,51). Integrating human, animal, and environmental surveillance data with population genomic analyses across geographic regions is critical to contain the threats posed by emergent *Salmonella* Infantis and its megaplasmids.

## Acknowledgments

Powered@NLHPC: This research was partially supported by the supercomputing infrastructure of the NLHPC (CCSS210001).

The authors of this study were supported by Agencia Nacional de Investigación y Desarrollo de Chile -ANID-through grants Fondecyt Postdoctorado Folio 3230796, Fondecyt Regular Folio 1231082, FONDEF IDeA I+D Folio ID24I10291; and scholarships Beca de Doctorado Nacional Folio 21241909, and Folio 21251661.

